# A new experimental evidence-weighted signaling pathway resource in FlyBase

**DOI:** 10.1101/2023.08.10.552786

**Authors:** Helen Attrill, Giulia Antonazzo, Joshua L. Goodman, Jim Thurmond, Victor B Strelets, Nicholas H. Brown, the FlyBase Consortium

## Abstract

Research in model organisms is central to the characterization of signaling pathways in multicellular organisms. Here, we present the systematic curation of 17 Drosophila signaling pathways using the Gene Ontology framework to establish a comprehensive and dynamic resource that has been incorporated into FlyBase, providing visualization and data integration tools to aid research projects. By restricting to experimental evidence reported in the research literature and quantifying the amount of such evidence for each gene in a pathway, we captured the landscape of empirical knowledge of signaling pathways in *Drosophila*.

**Summary statement:** Comprehensive curation of Drosophila signaling pathways and new visual displays of the pathways provides a new FlyBase resource for researchers, and new insights into signaling pathway architecture.

## Introduction

Signaling pathways are vital to life, allowing cells to respond to cues such as extracellular messengers, nutrients levels, tissue damage and infection, to maintain homeostatic control and to execute developmental programmes. From what are just a small number of developmental pathways, multiple cell and tissues types are generated and co-ordinated to form complex organs and systems (Pires-daSilva and Sommer, 2003). Research using *Drosophila* was central to the discovery and elucidation of many key signaling pathways, such as Notch, Hedgehog, Hippo and Toll, with the genetic approaches possible in *Drosophila* synergising with biochemical approaches utilizing vertebrate cells in culture. Using *Drosophila* to study pathways is no less popular today; searching the abstract/title of papers in PubMed with the terms ‘signaling’ and ‘Drosophila’ show the publication of >500 papers/year since the year 2000. Importantly, although the main ligands and receptors have been identified for each pathway, numerous regulators and effectors continue to be discovered.

Knowledge of signaling pathways can be recorded and integrated in a variety of ways. In review-style representations, signaling pathways are often presented as a sequential procession of events, simplified so that only the main components are shown. But signaling pathways are seldom, if ever, linear. They are complex, branching and can be regulated at multiple points and in context-dependent manners (Pires-daSilva and Sommer, 2003). Furthermore, it can be difficult to distinguish between gene products whose function is solely involved in a particular signaling pathway and those that are more promiscuous.

Online resources such as Reactome (Gillespie et al., 2022) and KEGG (Kanehisa and Goto, 2000) provide valuable bioinformatic summaries of pathways but are restricted to the main players. Given the volume of active research on pathways, it would be valuable to provide a resource that could be rapidly updated in response to new discoveries, including the numerous single observations that may not be followed up by subsequent studies, providing comprehensive coverage and allowing users to see that the evidence supporting some components is much stronger than for others.

We therefore set out to design a curation strategy that would capture the richness of the experimental research landscape on Drosophila signaling pathways, that could be maintained and scaled with ease. The widely used Gene Ontology (GO), a structured vocabulary used to annotate gene/gene product biological function (Ashburner et al., 2000; Gene Ontology Consortium, 2021), was used to assign genes as either core components or regulators of a particular pathway. An experimental-evidence weighted pathway resource was designed based on counting research papers with GO annotations supported by experimental evidence codes as a proxy for the strength of support for a gene’s involvement in a pathway. This data was assimilated into the Drosophila database, FlyBase (Gramates et al., 2022), in a way that can be easily sustained by future curators and presented with the aim of facilitating and seeding future research. Importantly, this resource displays the different pathways in the same format, highlighting the differences and similarities between these pathways. Comparison of pathway make-up reflects their deployment in response to stimuli and in development.

## Results and Discussion

### Curation of signaling pathways using annotation with the Gene Ontology to quantify experimental support

As a first step toward developing a signaling pathway resource for FlyBase, we sought to review and improve the curation of signaling pathway genes using the GO. The GO consists of a controlled vocabulary of specific biological terms linked in a hierarchical manner (Ashburner et al., 2000, Gene Ontology Consortium, 2023). Terms in the ‘signal transduction’ (GO:0007165) branch of GO biological process are used to annotate genes that encode members of pathways using the relationship ‘is_a’ (Fig. 1A). In the GO, processes may also have the relationship type ‘regulates’, and therefore genes are also labelled with terms that reflect this relationship to the pathway. Taking as an example the Epidermal Growth Factor Receptor (EGFR) signaling pathway, the receptor, Egfr, and the Erk mitogen-activated protein kinase, rolled, which are both essential for the execution of the pathway, are annotated with ‘epidermal growth factor receptor signaling pathway’ (GO:0007173), whereas argos, an antagonist of the pathway, is annotated with ‘negative regulation of epidermal growth factor receptor signaling pathway’ (GO:0042059). For some pathways, such as the EGFR pathway, very specific gene products are required for the biogenesis of pathway ligand, such as Rhomboid family proteases, required for the production of soluble, secreted EGFs (Urban et al., 2002). The biological role of such components can be annotated with terms that capture this e.g. epidermal growth factor receptor ligand maturation (GO:0038004). Thus, GO terms can be used as a handle to place pathway members and their regulators into simple categories, in our case to define a gene as either a core pathway member, a positive or negative regulator or involved in ligand production (Fig. 1A).

**Fig 1.**
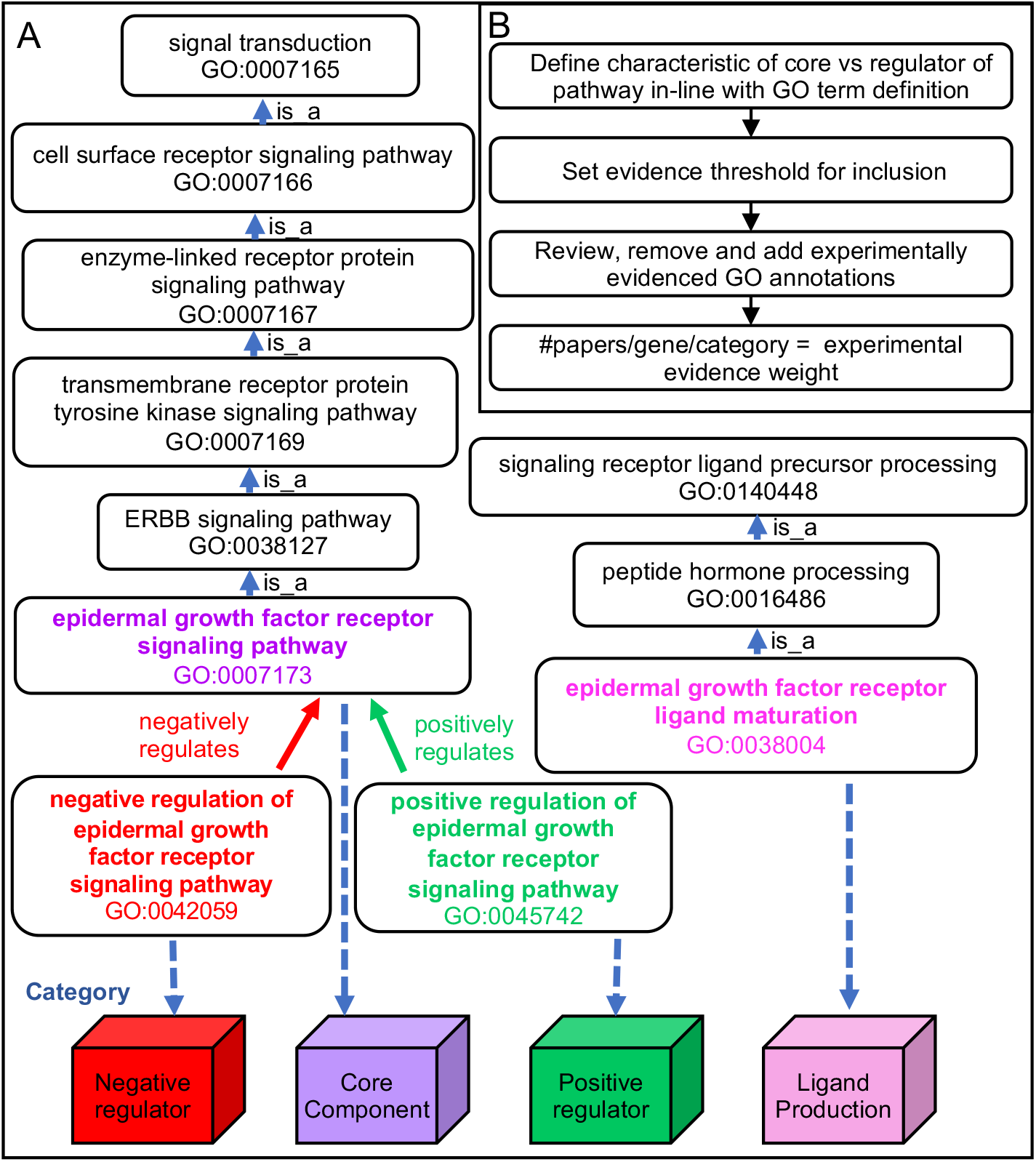
Strategy for experimental evidence-weighted curation of signaling pathways, using the EGFR signaling pathway as an example. (A) How use of GO to curate pathway research papers allows the terms to be used as labels to place pathway members in different categories. The structure of the GO relevant to the EGFR signaling pathway is used as an example. (B) Schematic of the signaling pathway review, curation process and calculation of evidence weight for inclusion.

A standard GO annotation links a GO term to a gene and includes an evidence code and the data source (e.g. publication) for that specific assertion. Evidence codes are used to distinguish the provenance of the assertion. For example, experimental evidence codes include ‘inferred from direct assay’ and ‘inferred from mutant phenotype’ and computational inference evidence codes include ‘inferred from electronic annotation’. Curating multiple examples of the same assertion from different research publications can be used to quantify the support for a given gene’s involvement in a pathway.

A set of 16 well-characterized signaling pathways were selected for review and curation. An important aspect of this review was developing a clear set of rules that can be consistently followed to curate the pathways (see materials and methods). Each pathway often has its own set of assays used as a readout of pathway activation, such as: expression of particular target genes (native or transgenic reporters e.g. 10xSTAT92E-luciferase for JAK-STAT (Chen et al., 2014); signature phenotype(s) (e.g. Border follicle cell migration for the PDGF/VEGF-receptor related (Pvr) pathway (Schober et al., 2005); or other readouts (e.g. the translocation of armadillo (β-catenin) into the nucleus for Wnt-TCF signaling (Tolwinski et al., 2003). Thus, although there is variation in the way pathways are studied, standards were established for a particular pathway and applied consistently during curation. Application of these rules also involved reviewing existing GO annotations and either validating them or removing them if incorrect. We then counted the number of papers associated with each gene annotated to a pathway that was supported by an experimental evidence code to give the ‘weighted’ evidence (Fig. 1B).

As an example, we describe how this protocol impacted on the genes included in the EGFR signaling pathway. At the start, 65 genes were already annotated to GO terms related to the pathway using both experimental and non-experimental evidence codes. Following our review of the literature, 32 genes removed that did not meet our criteria, 33 were retained, 47 were added (Fig. 2A-C; note that Fig. 2A reports gene-GO term associations, and since a gene may be associated with more than one GO term (e.g. Src42A is annotated with both positive and negative regulatory terms for EFGR signaling), the numbers shown are greater than the number of genes). Many of the genes removed contributed to processes that indirectly affect EGFR signaling, and were initially inferred to be part of the pathway based on experiments using a specific assay, where they phenocopied EGFR signaling mutants and/or had a genetic interaction with pathway members. For example, components of the multivesicular-body sorting-pathway ESCRT complexes, such as Vps28, Vps2 and Chmp1, when disrupted affect ligand-dependent internalization and destruction of Egfr, but as this is a separate biological process that acts on many proteins, upon review these genes were no longer considered to be specific to the signaling pathway itself (Fig. 2B). Genes meeting the criteria to be components of the EGFR pathway were mapped to one of four categories depending on their GO annotation (Fig. 1A, 2A). By a concerted effort to cover the research literature, we were able to capture many single studies where a gene was shown to have a role in the context of EGFR signaling. Curating each instance that met the annotation criteria and counting the number of research papers per gene per term, reveals the extent to which experimental research supports each gene’s role, thus building a picture of the experimental landscape of EGFR signaling research (Fig. 2D). Over half of the genes in the EGFR pathway group are supported by only one experimental observation, many of which are categorized as regulatory components. Our comprehensive identification of these single papers will help researchers find experimental results that may confirm their own observations and avoid duplication of effort.

**Fig 2.**
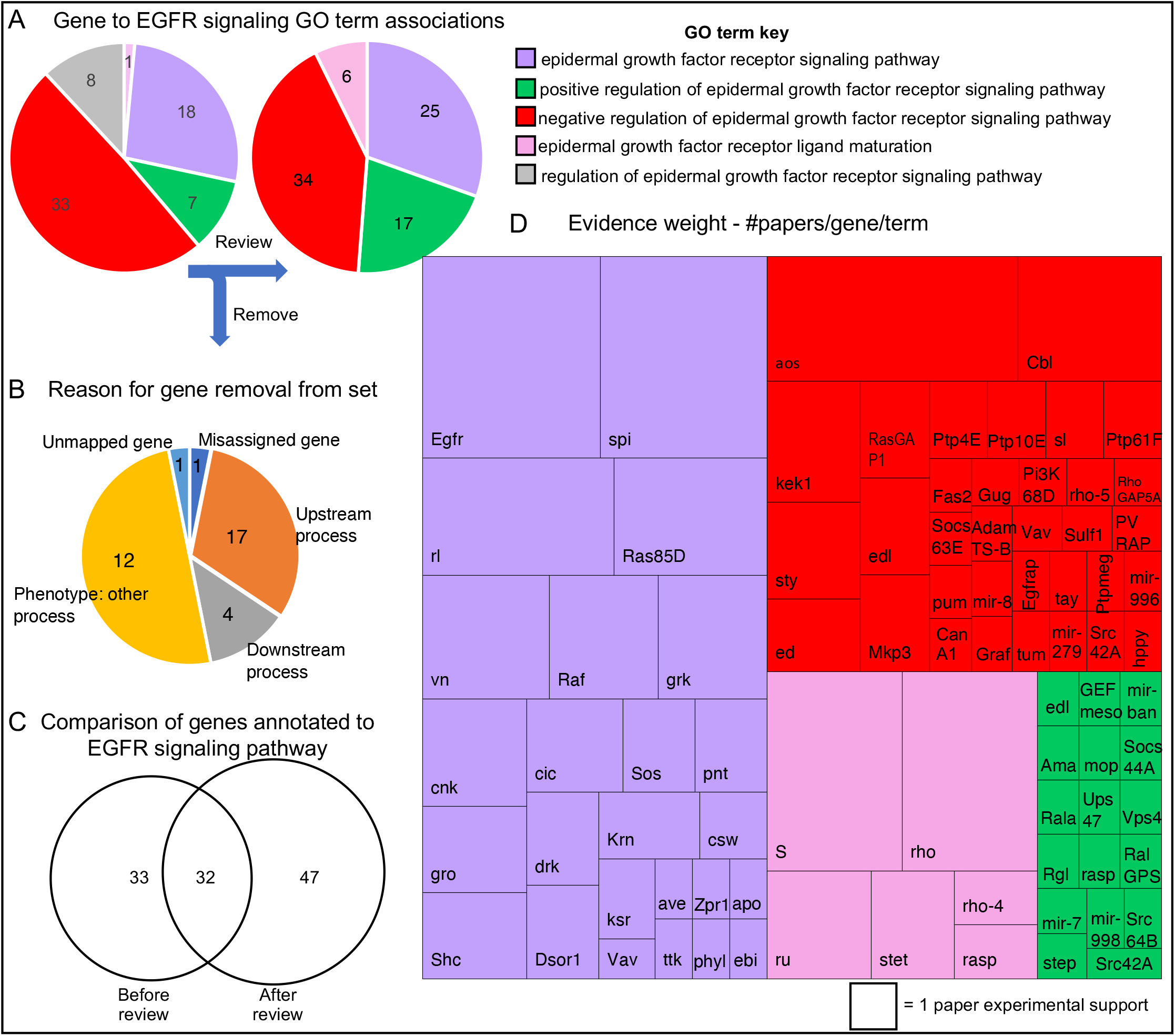
Evidence-weighted curation of the EGFR signaling pathway. (A) The number of genes associated with each EGFR-related GO terms before (left pie chart) and after review (right). The key is shown to the right. During the review process, all regulatory components were classed as positive or negative regulators and therefore there were no direct annotations to ‘regulation of epidermal growth factor receptor signaling pathway’ (grey segment) after review. (B) Pie chart displaying the reasons for removal for 32 genes previously annotated to the EGFR pathway, but which did not pass our curation criteria. (C) A Venn diagram GO annotated gene sets, before and after review, summarizes the extent of revision. (D) A treemap chart displaying experimental evidence weight. The block size is proportional to the number of research papers supporting each gene’s role in the EGFR signaling pathway. The corresponding gene symbols are shown. The term key is the same as for Fig. 2A.

### Comparison of the regulatory landscape of pathways from the results of experimental evidence-weighted curation

The number of genes associated with each pathway varies widely, from 13 (Activin) to 100 (Wnt-TCF) genes (Fig. 3). We wondered whether this reflected the inherent complexity of the pathway or was an indirect consequence of the length of time the pathway had been studied, i.e. more accumulated research. The correlation between the number of curated pathway members and the year a selected defining/prototypic member of the pathway (e.g. dome for JAK-STAT; dpp for BMP; sev for sevenless) was first described was only moderate (Supplementary Fig. S1; R^2^ = 0.40, p-value = 8.81×10^−3^). The four pathways with the largest number of members: Wnt-TCF (100 genes), Hedgehog (96), Notch (83) and Hippo (80), the date of the first publication on prototypic genes spanned nine decades: wg (1923), hh (1950), Notch (1916) and hpo (2002), respectively. Therefore, the number of components in each pathway appears to reflect some intrinsic features of the pathway, which may include the diversity of its roles and associated regulation, which have in turn contributed to extent of the literature for each pathway.

**Fig. 3.**
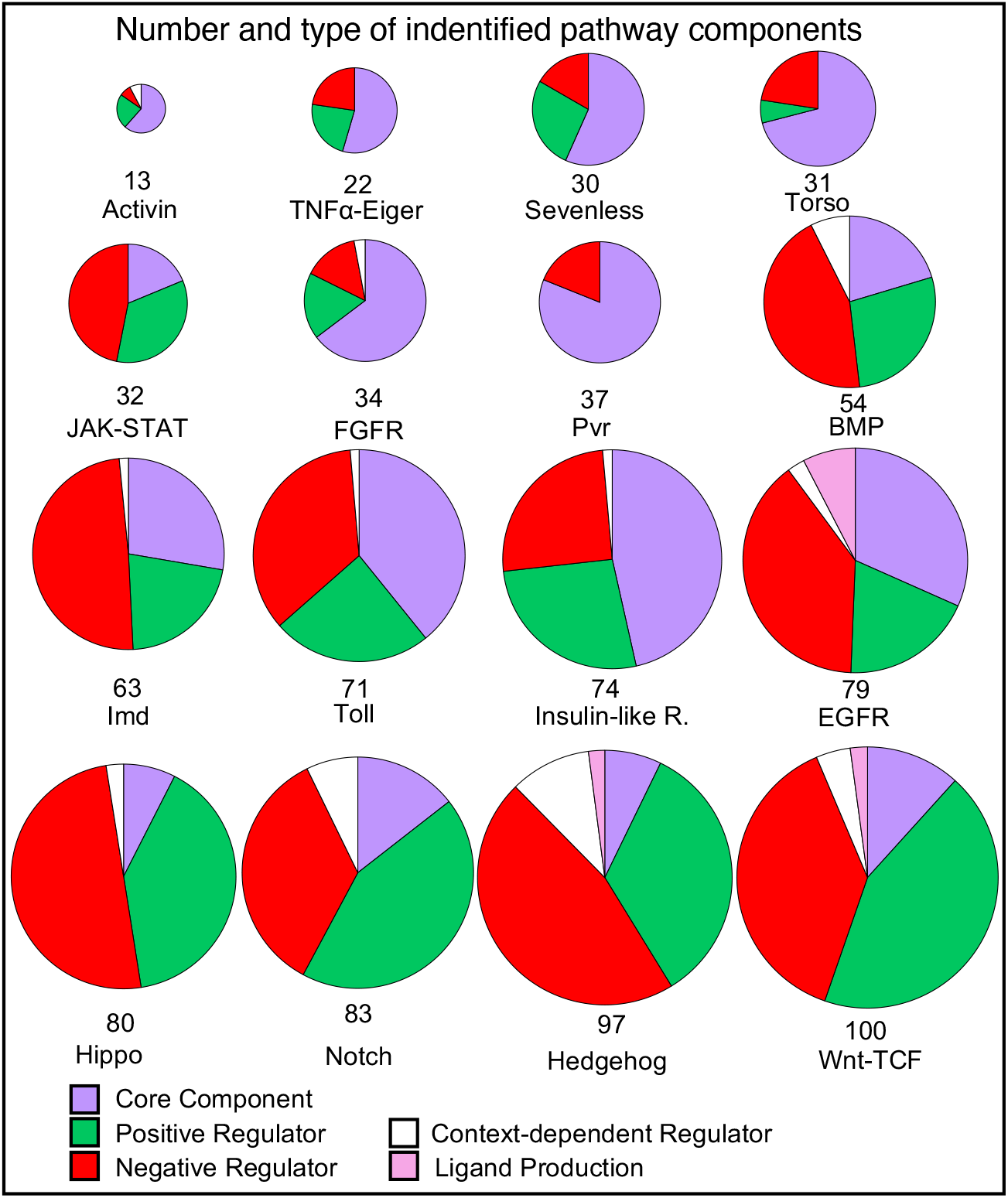
Distribution of types of components in each pathway. The number of genes associated by GO annotations for each pathway are illustrated as pie charts. Each pie chart is broken down into the categories indicated by the key. The total number of genes in the pathway is illustrated by the diameter of the circle and the number at base. The pathways are ordered by the number of component genes. Where genes have different regulatory effects in different contexts, they are classed as context-dependent regulators. Note that for this figure, the Toll signaling pathway, which has a more complex extracellular activation cascade, those genes annotated to ‘negative regulation of Toll receptor ligand protein activation cascade’ (GO:0160035) are included in the ‘Negative Regulator’ category, and genes annotated to ‘Toll receptor ligand protein activation cascade’ (GO:0160032) are included in the ‘Core Component’ category.

Looking at the classification of pathway components, for most of the pathways (10/16) the number of regulatory components exceeds the number of core components (Fig. 3) and for six pathways (JAK-STAT, BMP, Hippo, Notch, Hedgehog, Wnt-TCF) over 80% of the components are regulators. A general observation is that pathways that are used most frequently in development and those that are involved with immune processes are more heavily weighted towards regulators. This is elaborated on with examples below.

The innate immunity pathways - JAK-STAT, Toll and Imd, display a high proportion of negative regulators (37-49% of components). Unchecked immune response can lead to damaging inflammation, as such down-regulation of these pathways after activation is a vital part of their circuitry. For example, both the Toll and Imd pathways can be negatively regulated by the removal of extracellular stimuli. For Toll, the Serpins (SERine Protease INhibitors) dampen the ligand-activating zymogen cascade (Fullaondo et al., 2011) and for the Imd pathway, bacterial peptidoglycans are removed by enzymatically active peptidoglycan recognition proteins (Costechareyre et al., 2016).

In *Drosophila* the Hedgehog, Wnt-TCF, Notch, BMP and EGFR pathways are repeatedly used in development in numerous contexts. The portion of regulators curated for these pathways is high (91-61%). First, as these pathways are used in so many different developmental contexts, it would be reasonable to assume that the range regulators encountered, in terms of context such as tissue- and temporal-specific or precision control needed for patterning, would be correspondingly diverse. An example of the complex interplay in context-dependent control is that of the numb and sanpodo, that only regulate Notch signaling during asymmetric cell division. In the absence of numb, sanpodo positively regulates Notch signaling but in the presence of numb, sanpodo inhibits Notch signaling.

Numb, which is inhibitory, can only act in the presence of sanpodo (Babaoglan et al., 2009). Thus, the asymmetric distribution of numb drives acquisition different cell fates. Second, the strict control of a signaling window is essential for the execution of developmental programs - when a pathway is switched off is as important as when it is switched on (Perrimon et al., 2012). Some pathways need to be continuously repressed in the absence of signal, notably Wnt-TCF (Roberts et al., 2012) and Hedgehog pathways (Han et al., 2015), which are de-repressed by the presence of ligand. Pathways such as receptor tyrosine kinase (RTK) pathways are principally controlled by the availability of receptor ligands rather than having to be actively repressed. The EGFR pathway has a high proportion of negative regulators, 39% of components, in comparison to other RTK pathways, which most likely represents its widespread usage in development and has a similar regulatory profile to BMP signaling, which has 44% negative regulators (Shilo, 2005). Thus precise spatial and temporal control is required to build complex systems and structures.

Comparing the RTK pathways in order of number of members: Torso, Sevenless, FGFR, Pvr, Insulin-like receptor, EGFR (from 30 to 79 genes), reflects the scope of their use. Of the RTK pathways, the Torso and Sevenless have smallest number of components (31 and 30 genes, respectively). Both pathways are restricted in their use - Torso signaling is principally involved in the development of embryonic termini and Sevenless for the specification of the R7 photoreceptor cells. The lack of complexity of these pathway reflects their extremely limited developmental deployment. The Pvr pathway is an interesting contrast in having a large proportion of genes (81%) characterized as core components. Of all signaling pathway curated, the Pvr appears to be the most branched, capable of employing several intracellular cassettes such as the Erk kinase cascade, PI3K/TORC1, Rac/Rho signaling and the JNK cascade. It is associated with inducing a number of processes, such as border follicle cell migration, epidermal wound healing, hemocyte spreading and cell proliferation (Tsai et al., 2022, Mues et al., 2023).

In contrast to the Pvr pathway, the Hippo pathway has a particularly high proportion of regulators (92%), which may reflect the substantial difference in logic of this pathway. It is an intracellular cassette in which hippo kinase phosphorylates warts kinase which in turns phosphorylates the transcriptional coactivator yorkie leading to its cytoplasmic retention (Huang et al., 2005). The regulatory components include many membrane proteins and scaffolding proteins, as the hippo pathway integrates mechanical cues to manage cell growth and survival and its balance is key in determining organ size and tumour suppression (Chang et al., 2019). Pathway components are therefore heavily weighted towards regulation (Pires-daSilva and Sommer, 2003).

### Developing a research-based pathway resource

The next step was to develop a signaling pathway resource within FlyBase to facilitate future research. Pathway Reports were generated using a similar architecture to the FlyBase Gene Group resource (detailed in Attrill et al., 2016), with key differences to maximize usefulness to pathway research. Pathway Reports are organized in a hierarchical fashion, with a top-level parent report and sub-groups, and we divided pathway components into the sub-groups of ‘core members’, ‘positive regulators’, ‘negative regulators’ and, in some cases, specific ‘ligand production’ sub-groups, mirroring the curation scheme in Fig. 1A. The reports can be queried using the FlyBase QuickSearch ‘Pathways’ tab on the homepage (http://flybase.org/) and a list is available at http://flybase.org/lists/FBgg/pathways. As described more fully below, the pathway reports contain: 1) a customizable table of pathway members; 2) a visual summary of the GO annotation (the GO ribbon stack); 3) two types of pathway diagrams, static thumbnails and dynamic physical interaction networks; and 4) links to other resources.

#### A customisable table of pathway members as a resource and information hub

Each gene belonging to a pathway sub-group is represented as a row in the table, and by default the columns contain the following data: Gene Symbol; Gene name; Gene Group Membership; GO Molecular Function (Experimental); and # Pathway references (Supplementary Fig. S2). Additional columns can be selected: # All Research Refs; Also Known As; Antibody; Classical / Insertion Alleles; Transgenic constructs; Disease Models (Experimental); Potential Disease Models; Other Pathways; and Human orthologs. The user can select which columns are: shown or hidden; order based on the column value; or filter the entries by text match. The ‘# Pathway Refs’ column contains the number of GO-curated papers that provide direct experimental evidence for a gene’s role in a pathway. Clicking the hyperlinked number in this column generates a FlyBase ‘HitList’ of these references, which can be downloaded or analysed further. Thus, the user has access to the molecular function of each gene product, the literature, the experimental reagents used in the study of each gene and links to human disease and fly models of those diseases.

#### GO ribbon stack

To enable users to compare the biological functions of the pathway components, we chose to display a stack of GO summary ribbons (Fig. 4A). These summaries take advantage of the hierarchical nature of GO to group the detailed GO annotations into broad categories, each represented as a square within the ribbon; the presence of GO annotations in each category colors them, from faint to dark blue, depending on the number of annotations. Clicking on each colored square displays more detail, as indicated in Fig. 4A. In this way, a user can easily get an idea of which genes encode enzymes or extracellular proteins and whether the pathway has more prominent roles in, for example, the immune, nervous or reproductive systems and quicky see the more detailed information.

**Fig. 4.**
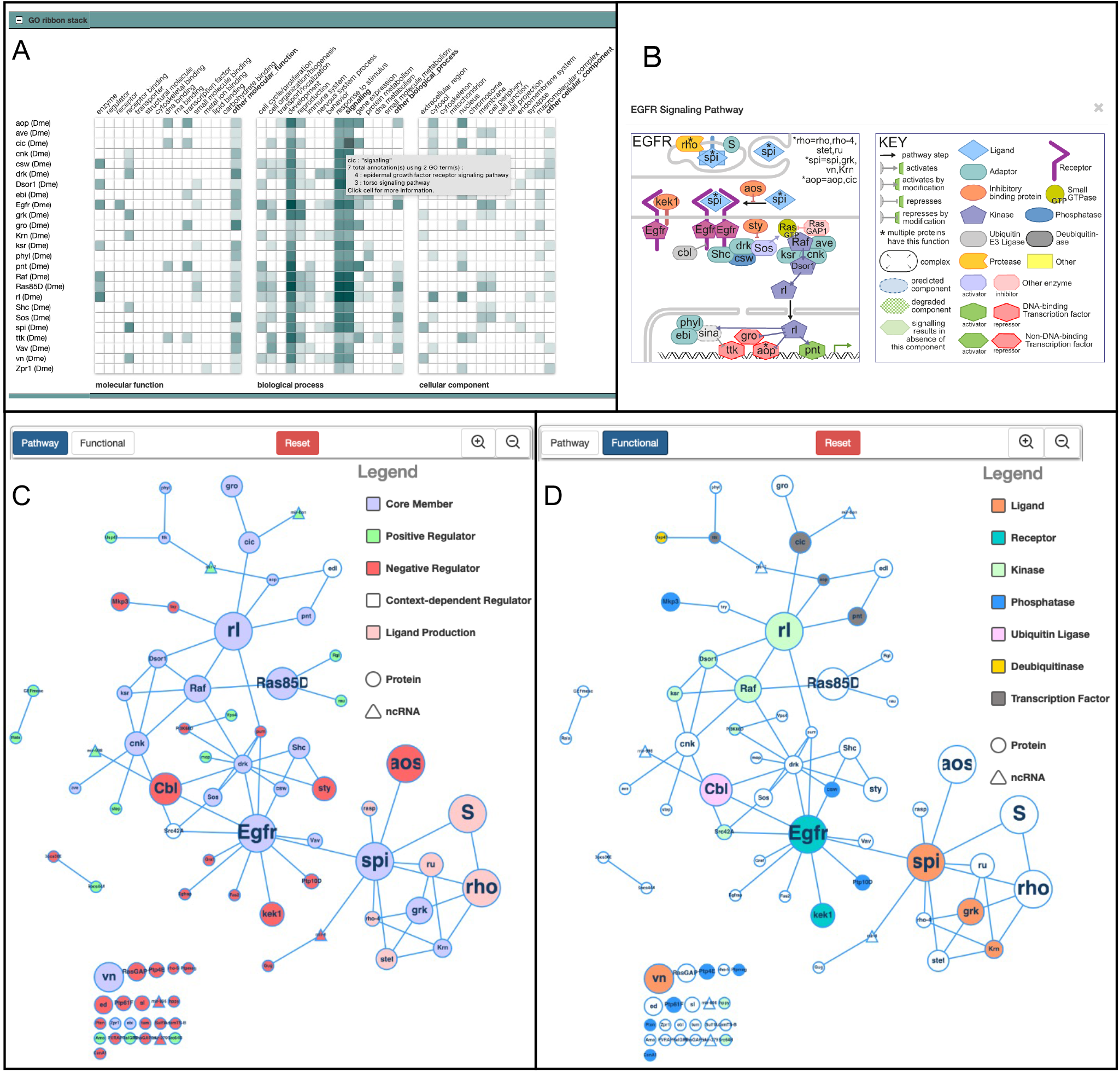
FlyBase Pathway Report Pages. Three visual summaries are presented on the Pathway pages, exemplified with the EGFR Signaling Pathway. (A) The GO ribbon stack is a graphical summary of the GO annotations for each gene, with annotations grouped under high-level categories. The color intensity of each cell indicates how many unique terms are grouped in that particular category. The unique terms that are grouped under a particular cell can be seen by hovering over or clicking on the cell, as shown for one of the cells of *cic*. (B) A thumbnail review-style image for the EGFR Signaling Pathway. (C, D) A physical interaction network in which the nodes represent genes, the sizes of which are proportional to the number of supporting papers, up to ≥10. (C) Shows the ‘Pathway view’ and (D) the ‘Functional view’ (for visual simplicity, we selected fewer higher level categories compared to the thumbnails). The edges represent physical interactions between components, from FlyBase. Unconnected nodes are also displayed.

#### Pathway diagrams

To display a visual summary of each pathway, two contrasting approaches were developed. To give a relatively simply summary of how each pathway works, a static ‘Thumbnail’ for each pathway is provided, using a standard review-type representation that shows the position and role of the components in the cell, and gives a sense of the route of the pathway from outside the cell into the nucleus (an example, the EGFR signaling pathway thumbnail, is shown in Fig. 4B and all thumbnails are shown in supplementary Fig. S3). The Thumbnails focus on the core pathway members, so they did not become too complex and consulted with experts on each pathway and reviewed the curated evidence weight to make sure we had included the key players (see acknowledgements). By using a uniform format for depicting the different types of molecules, the molecule architecture of the different pathways can be easily compared. The Thumbnails are available within FlyBase as downloadable SVG images, so they can be modified by researchers if desired. They are displayed next to a concise textual summary of the pathway.

We also sought a dynamic way of visualising all the pathway members with information about their function and the weight of evidence for their inclusion in the pathway, generated computationally from the data in FlyBase so it would be automatically updated to include new data. For this, each pathway member is displayed as a node in a network of physical interactions (Fig. 4C and D), generated with Cytoscape.js (Franz et al., 2016). The size of each node is a function of the number of curated papers that report experimental evidence for the membership of the gene in the pathway (with a maximum size at ≥10 papers). The edges between nodes are derived from physical interaction data curated in FlyBase. Post-transcriptional regulation by non-coding RNAs (triangular nodes) are integrated into the interaction network to show where regulatory intersections may occur.

Users can switch between two color-coded schemes: a ‘Pathway’ view (Fig. 4C), where the nodes are colored according to high-levels roles in the pathway (Core Member; Positive Regulator; Negative Regulator; Context-dependent Regulator; Ligand Production); and a ‘Functional’ view, with the node color corresponding to selected molecular function classes (Fig. 4D). The networks can be rearranged by users by manually dragging nodes, which will maintain their position between views, or by resetting the view. As these networks are updated each FlyBase release through in-house computational pipelines, they reflect the current state of curated knowledge.

#### Links to other resources

The Pathway Reports are designed to act as a knowledge hub. The customizable table provides links to other data and resources in FlyBase. In addition, links are provided to relevant FlyBase Gene Groups e.g. links from the ‘Wnt-TCF Signaling Pathway Core Components’ page to the ‘FRIZZLED-TYPE RECEPTORS’, ‘WNTs’, ‘WNT ENHANCEOSOME’ Gene Groups pages. In the External Data section, linkouts are provided to external resources, such as Reactome, KEGG, Wikipathways and The Interactive Fly.

#### A scalable model - Adding a newly described antiviral pathway

One of the core strengths of this pathway resource is that the underlying model is scalable and easy to maintain. A common addition to our set of well-characterized pathways is the description of new regulatory interactions. However, we also intend to grow this resource with more signaling pathways – a recent example of which is the addition of the cGAS/STING Signaling Pathway. In vertebrates, STING (stimulator of interferon genes) was first shown as a component of innate immunity, involved in the induction of interferon (Ishikawa and Barber, 2008; Ishikawa and Barber, 2008). STING is activated by cyclic GMP-AMP (cGAMP), which is produced by cGAMP synthase (cGAS) in response to cytosolic DNA, e.g. viral DNA. STING then activates IKK and TBK1 and, further downstream, the activation of the transcription factors IRF3 and NF-κB leads to the transcription of immune response genes. In insects, although *Sting* had been linked to defence response (Goto et al., 2018; Martin et al., 2018) and NF-κB (Rel) activation, the cGAS component remained elusive until Slavik et al. (2021) and Holleufer et al. (2021) identified two cytosolic cGAS-like gene that respond to viral infection. It was therefore apparent that the cGAS-STING module was an evolutionary conserved pathway with the potential to respond to other pathogen-associated molecular patterns such as dsRNA. As such, a new GO term ‘cGAS/STING signaling pathway’ (GO:0140896) was created in 2022, allowing pathway components to be specifically annotated to this pathway rather than to a more general terms related to ‘defense response to virus’ (GO:0051607) or canonical NF-kappaB signal transduction (GO:0007249). Given the relative novelty of this pathway, so far only 6 core components have been annotated to it. FlyBase curators will continue to populate this page as more experimental information arises, keeping pace with research. The importance of this pathway is highlighted as *Drosophila* grows as a popular model for virus infection, including viruses with a direct impact on human health such as EBV, HIV-1, SARS-CoV-1 and -2, Zika, dengue and West Nile fever viruses (Hughes et al., 2012). The addition of the cGAS/STING pathway brings the total number of pathways in the resource to 17.

## Conclusions

This paper describes a novel experimental-evidence weighted approach to assembling a signaling pathway resource. Using GO curation to redundantly curate the same experimental conclusion from multiple research papers provides a mechanism for measuring the strength of characterization of a gene to pathway relationship. By presenting this information in a tabular and graphical way in dedicated pages in FlyBase, researchers can easily distinguish ‘central players’ from single observations identifying new components. From a curatorial perspective, this is easy to update, maintain and scale beyond the initial focused effort. As GO is machine-readable, widely used and distributed with many existing tools, this work adheres to FAIR (Findability, Accessibility, Interoperability, and Reusability) principles (Wilkinson et al., 2016). The pathway knowledge we have curated has already been incorporated by two bioinformatic projects: FlyPhoneDB, a tool used to predict cell-cell communication between cell types from Drosophila single-cell RNA sequencing data (Liu et al., 2022), and PANGEA, a Gene Set Enrichment Tool (Hu et al., 2023), demonstrating the utility of the resource beyond FlyBase.

The approach to curating and presenting information described in this paper is applicable across organisms and, as it is based on the GO, to any other term in the biological process branch. Integrating the GO curation of pathways into dedicated pages in FlyBase allows users to see other data side-by-side and make their own observations. An example of this is the link between genes annotated to ‘BMP signaling pathway’ (GO:0030509) and disease models for ‘juvenile polyposis syndrome’ (DOID:0050787) (Supplementary Fig. S4). To further improve this integrated system, we aim to impart directionality to some of the edges in the pathway networks by annotating which gene products are the targets in various interactions.

By a clear and simple division of components into core, regulatory and, in some cases, ligand production, we have been able to observe differences in the diversity and scale of regulatory interactions between pathways deployed in different contexts. An important aspect of our curation effort has been the inclusion of post-transcriptional regulation by ncRNAs - an element that is missed by other pathway modelling approaches. In *Drosophila*, approximately 2.7% of genes (485/17,896 mapped genes) encode miRNAs, which play a crucial role in the regulation of cellular processes by directing the activity of the RNA-induced silencing complex. In this project we have associated 31 unique miRNA genes with pathways. Some miRNAs regulate multiple pathways – for example, mir-8, mir-ban, mir-279, mir-958 and mir-7 have curated interactions with 3 or more pathways.

Most pathway resources must be, by pragmatic design, limited to an orthodox set of genes, thereby missing new and incremental knowledge. Repetition is often seen as the gold-standard for measuring confidence in a result. The evidence-weighted model for curation presented here allows a measure of reproducibility i.e. how often the observation has been made by separate studies. We believe that the dynamic model of pathway curation we have outlined here and its integration with other data curation streams, shows how a simple curation framework can be used produce a rich, up-to-date resource which can further feed back into research.

## Materials and methods

### Curation Strategy

For each pathway, an initial list of genes to review was downloaded from FlyBase using the FlyBase Vocabularies term tool queried with the appropriate pathway term (including child and regulatory terms). In concert with current reviews, papers and the relevant GO term descriptions, we established the definitions of what constitutes the core pathway boundaries. The initial list of genes was reviewed and annotations that did not fit the curation criteria were removed/replaced. The annotations for each pathway were made consistently at an appropriate level – for example, with genes annotated to ‘Wnt signaling pathway’ (GO:0016055), we replaced the term with ‘canonical Wnt signaling pathway’ (GO:0060070) where this was appropriate and this term was used as the handle to compile the list of Wnt-TCF Signaling Pathway Core Components. For this initial pass, we also focused on identifying research papers reporting experimental evidence for well-established pathway members that did not have any such evidence annotated to them. Further to this, we reviewed early, key papers that demonstrated the involvement of certain genes in pathways, including papers which had been previously missed as they pre-dated the introduction of the GO. As we reviewed papers, we established curation benchmark standards for pathway membership inclusion and recorded these as a Drosophila pathway curation guide (https://wiki.flybase.org/mediawiki/images/9/9f/FB_Pathway_Curation_Manual.pdf). Many pathways, especially those with extensive characterization, were subject to further rounds of curation after the establishment of pathway pages in FlyBase. Pathway pages were constructed by using the GO terms as markers of the inclusion of papers in a Signaling Pathway group (see Supplementary Table 1 for mapping).

### Construction of networks

The pathway networks are generated for every FlyBase release using Cytoscape.js (Franz et al., 2016) and an in-house pipeline that takes as input the FlyBase physical interaction file in MITAB format (Perfetto et al., 2019) and an internal FlyBase gene group membership reporting file, to produce a JavaScript file per pathway. The layout of the networks is established using the fCoSE algorithm (Genc and Dogrusoz, 2016; Balci and Dogrusoz, 2022), with the following parameters: “directed: true”, “nodeRepulsion: 450000”, “idealEdgeLength: 100”, “gravity: -500”. The networks are then integrated into the Cytoscape.js widget developed by FlyBase, available at https://github.com/FlyBase/cytoscape-widget.

### GO stacked Ribbons

The GO annotation ribbon stacks are generated using the GO Consortium Web Component (available at *github.com/geneontology/wc-ribbon)* with some in-house modifications and configuration, with input a FlyBase-generated file of gene IDs and FlyBase GO annotation data. The FlyBase GO annotation ribbon mapping categories used can be found at https://wiki.flybase.org/wiki/FlyBase:Gene_Report#Function and are labelled as subset: goslim_flybase_ribbon in downloads of the GO (http://geneontology.org/docs/download-ontology/). These FlyBase data are publicly available via the FlyBase API at http://flybase.org/cgi-bin/getRibbonJSON_agr-like.pl, with a “subject” parameter consisting of a list of FlyBase IDs. (Please note that this API address may change in the future, but any changes will be documented at https://flybase.github.io/api/swagger-ui/, in the FlyBase GitHub repository.)

### Version information

The data used was from FlyBase release FB_2022_06, GO version 2022-09-19. For comparison of before review, data from release FB_2017_02 was used.

## Supporting information

Table S1. Summary of Signaling Pathways content in FlyBase.

## Acknowledgments

We would like to thank David Strutt (University of Sheffield, UIK), Julius Mieszczanek (MRC Laboratory of Molecular Biology, Cambridge, UK), Alfonso Martinez-Arias (University of Cambridge, UK), Golnar Kolahgar (University of Cambridge, UK) and Nic Tapon (Francis Crick Institute, UK) for advice on the design of the Signaling Pathway pages. For reviewing the Pathway thumbnails, we would like to thank Erika Bach (New York University, US), Sarah Bray (University of Cambridge, UK), Edan Foley (University of Alberta, Canada), Marc Furriols (Institute of Molecular Biology of Barcelona, Spain), Jin Jiang (University of Texas Southwestern Medical Center at Dallas, US), Bruno Lemaitre (EPFL-GHI-SV-UPLEM, Switzerland), Steven Marygold (FlyBase, University of Cambridge, UK), Ian Mcgough (Babraham Institute, UK), Gillian Millburn (FlyBase, University of Cambridge, UK), Arno Müller, Universität Kassel, Germany, Mike O’Connor (University of Minnesota US), David Stein (University of Texas at Austin, US), Neal Silverman (UMass Chan Medical School, US), Hugo Stocker (Institut für Biochemie, Switzerland), Nic Tapon (Francis Crick Institute, UK), Jean-Paul Vincent (Francis Crick Institute, UK), Lei Xue (Tongji University, China). We also wish to thank members of the GOC and the Alliance of Genome Resources for useful discussion and contribution to signaling pathway guidelines, in particular Pascale Gaudet (Swiss Institute of Bioinformatics, Switzerland), Ruth Lovering (UCL, UK), Paul Thomas (University of Southern California, US), Kimberly van Auken (Caltech, US), David Hill (Jackson Laboratory, US) and, for the development of and help with implementing stacked GO ribbons, Laurent-Philippe Albou (Université Louis Pasteur, France).

Members of the FlyBase Consortium: Norbert Perrimon (PI), Nicholas H. Brown (Co-PI), Brian Calvi (Co-PI), Richard Cripps (Co-PI), Thomas Kaufman (Co-PI), Susan Russo Gelbart (PD), Giulia Antonazzo, Helen Attrill, Kris Broll, Seth Campbell, Lynn Crosby, Gil dos Santos, Kathleen Falls, Josh Goodman, Damien Goutte-Gattat, L. Sian Gramates, Victoria Jenkins, Pravija Krishna, Aoife Larkin, Ian Longden, TyAnna Lovato, Steven Marygold, Beverley Matthews, Alex McLachlan, Gillian Millburn, Arzu Ozturk-Colak, Clare Pilgrim, Jolene Seme, Victor Strelets, Christopher J. Tabone, Jim Thurmond, Vitor Trovisco, Pinglei Zhou and Mark Zytkovicz.

## Data availability

Data used for this analysis is available at DOI:10.5281/zenodo.7102381. This comprises the data files: physical_interactions_mitab_fb_2023_01.tsv.gz, pathway_group_data_fb_2023_01.tsv.gz and gene_group_data_fb_2023_01.tsv.gz. The current list of FlyBase Pathway Reports is available at http://flybase.org/lists/FBgg/pathways.

## Supplementary Material

**Legends**

**Table S1. Summary of Signaling Pathways content in FlyBase**.

An Excel spreadsheet summary of the Signaling Pathway page content in FlyBase release version FB_2022_06. The name and unique FlyBase Gene Group ID (FBgg#) are given in the first column. The second column is the GO term and ID that is mapped to the FBgg. The number of genes and papers linked to the pathway are given into the last two columns. Note that the sum of the numbers for the member groups is often different than for the top-level group as papers and genes may be shared between more than 1 subgroup. For each pathway group, except cGAS/STING, the table also includes the gene selected as the prototypic representative and year of publication of first research paper on the prototype.

**Figure S1.**
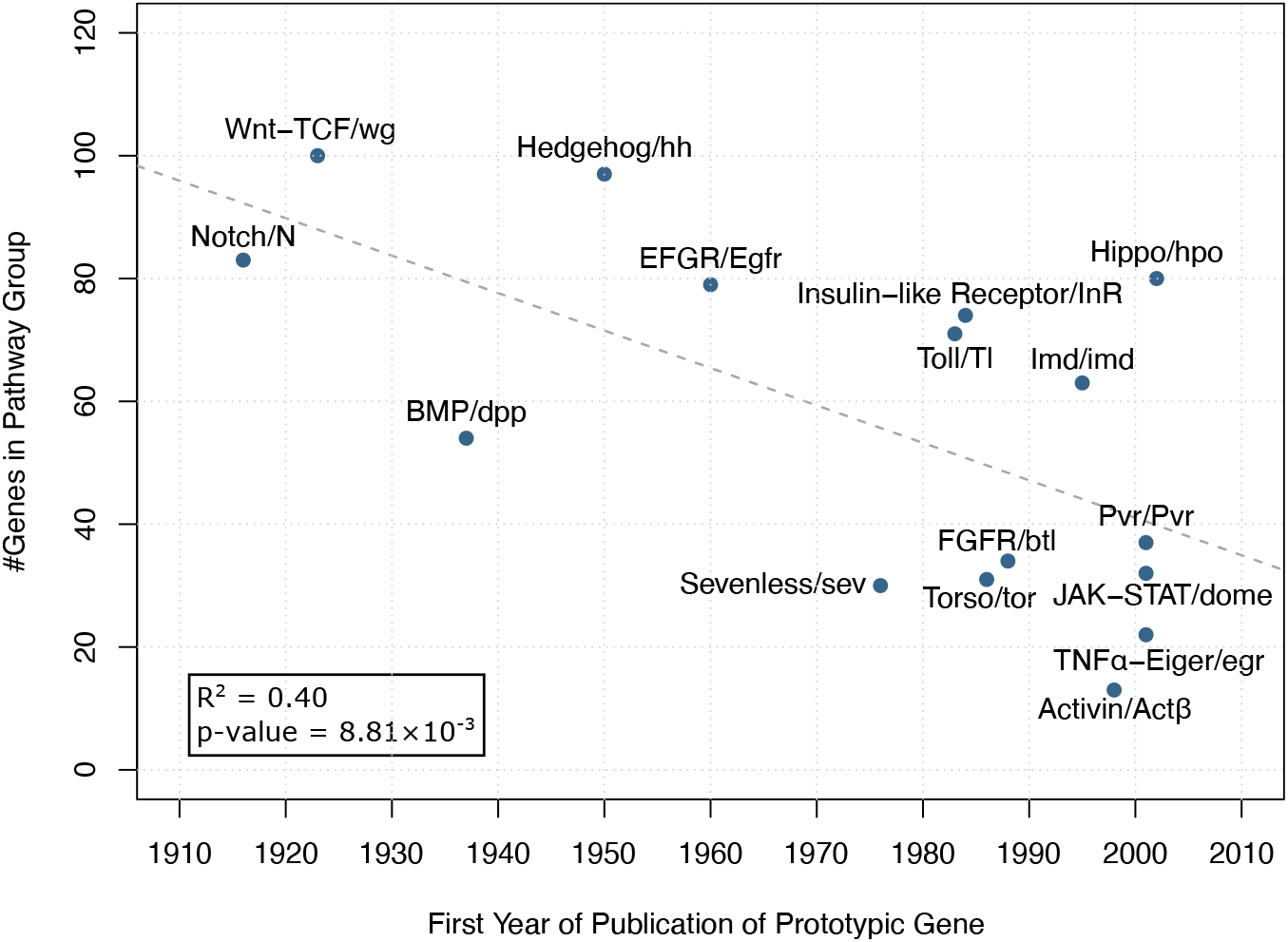
Pathways studied for a greater length of time do not necessarily have more identified components. The graph plots the year the prototypic member was first characterized, as measured by the first research paper associated with the gene in FlyBase, versus the number of curated pathway members. Linear regression was performed using the *lm* function in R.

**Figure S2.**
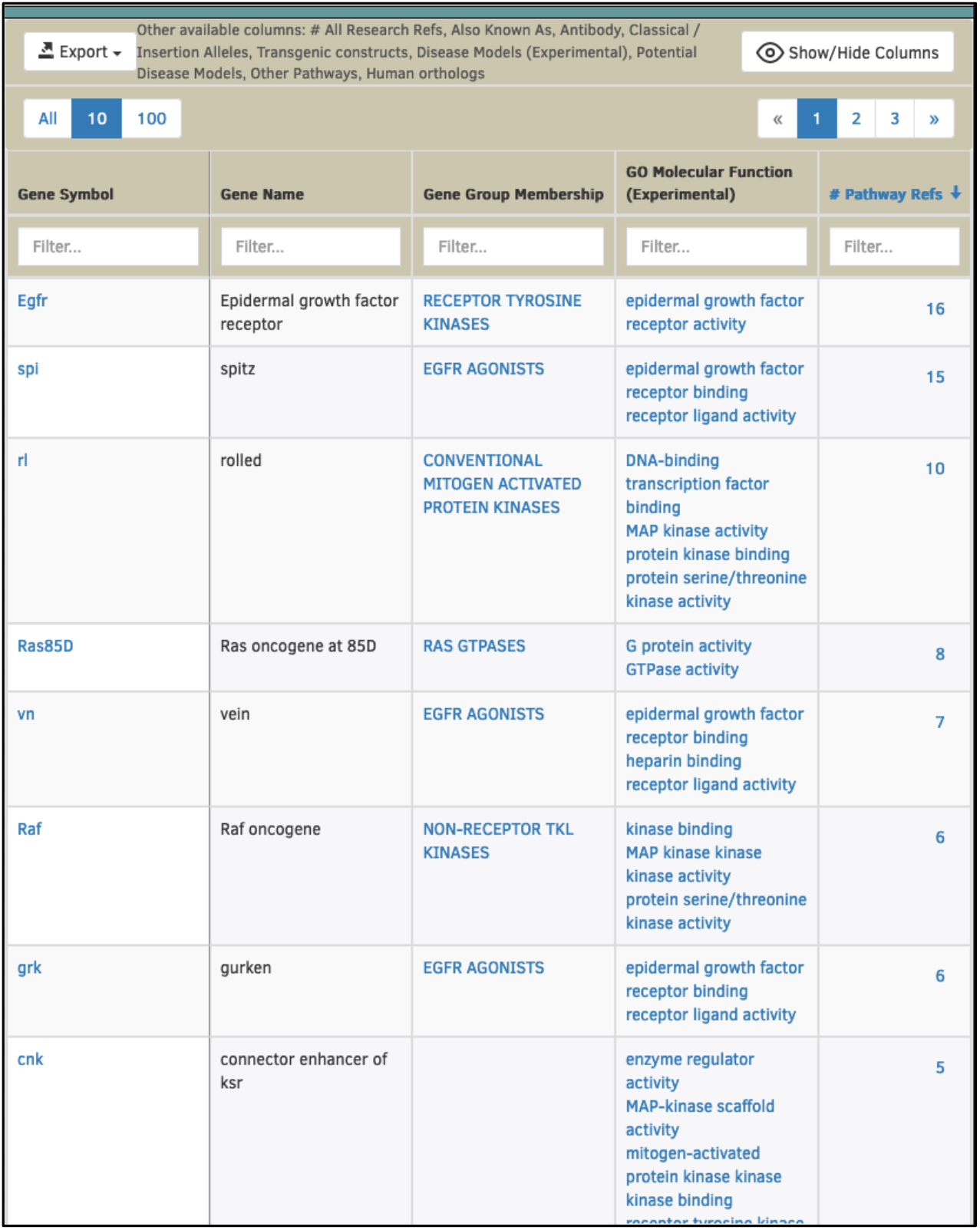
EGFR Signaling Pathway Core Components Members Table. A screenshot of the EGFR Signaling Pathway Core Components (FBgg0000951) members table showing the default column view sorted by # pathway refs (i.e. supporting papers for that gene annotated with ‘epidermal growth factor receptor signaling pathway’ (GO:0007173) using an experimental evidence code).

**Figure S3.**
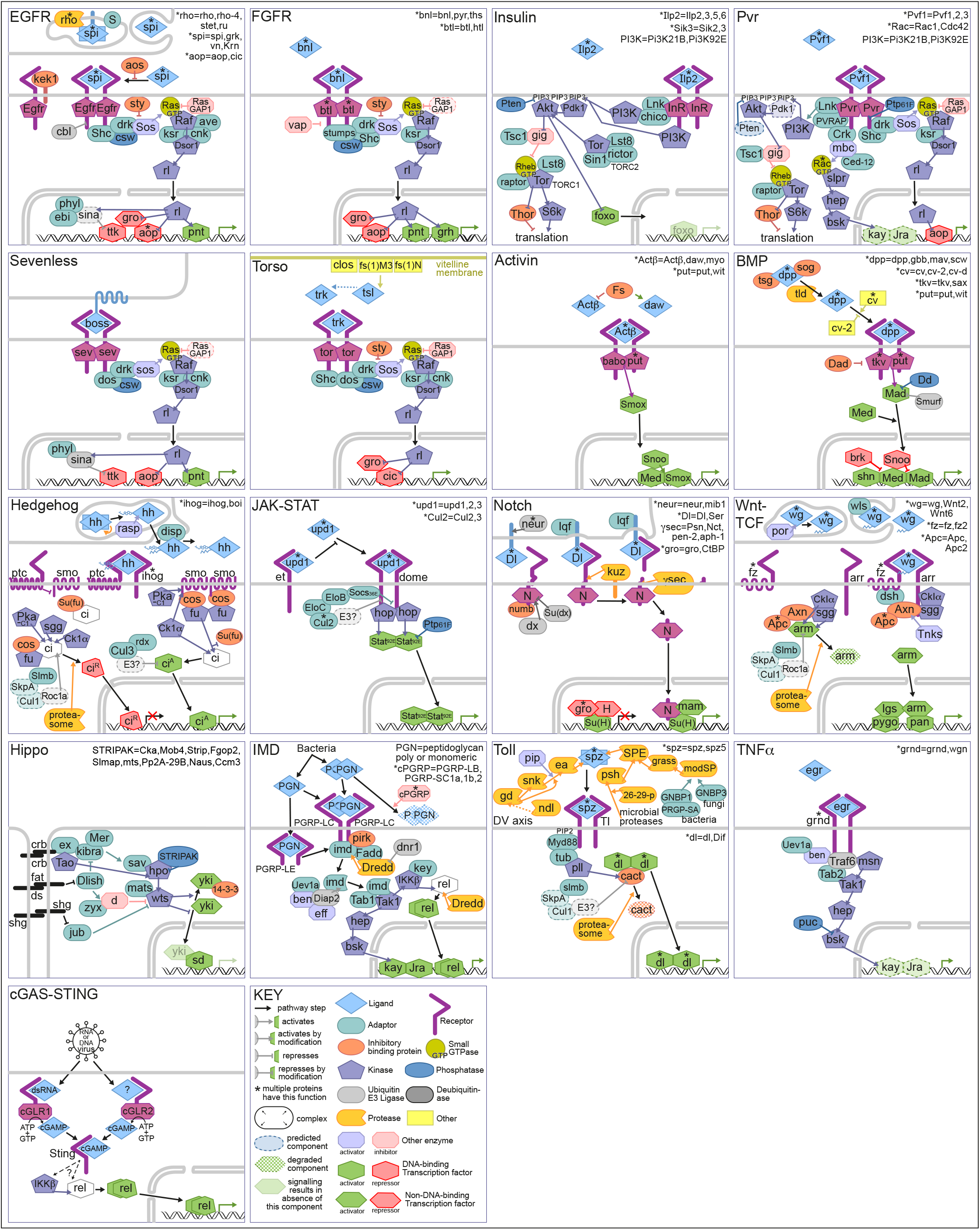
Thumbnails of signaling pathways in the FlyBase resource. The 17 pathways that have been reviewed by evidence-weighted curation are presented in FlyBase. For each, a text-book style over-view is available. These Thumbnail images were created in Adobe illustrator. The Pathway thumbnails were reviewed by experts in the field (see Acknowledgments).

**Figure S4.**
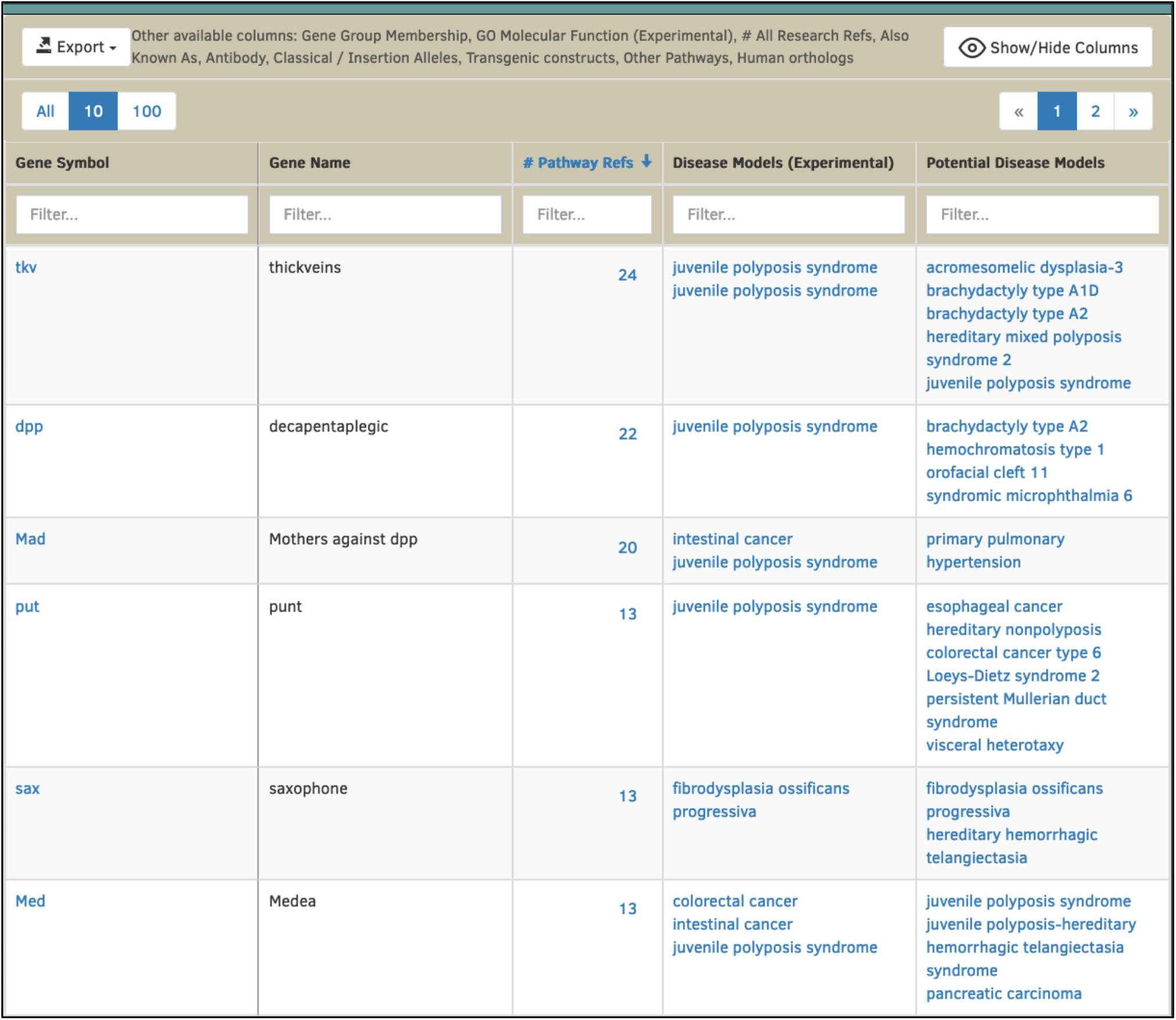
Disease associations for members of the BMP Signaling Pathway Core Components page. Columns showing experimental and predicted (computed from human-ortholog OMIM links) show that six members of the BMP Signaling Pathway Core Components (FBgg0001085) have been used to model ‘juvenile polyposis syndrome’ (DOID:0050787) in flies and two members have human orthologs with ‘juvenile polyposis syndrome’ associations in OMIM.

